# Cyclin E expression is associated with high levels of replication stress in triple-negative breast cancer

**DOI:** 10.1101/532283

**Authors:** Sergi Guerrero Llobet, Bert van der Vegt, Evelien Jongeneel, Rico D. Bense, Carolien P. Schröder, Marieke Everts, Geertruida H. de Bock, Marcel A.T.M. van Vugt

## Abstract

Replication stress entails the improper progression of DNA replication. In cancer cells, including breast cancer cells, an important cause of replication stress is oncogene activation. Importantly, tumors with high levels of replication stress may have different clinical behavior, and high levels of replication stress appear to be a vulnerability of cancer cells, which may be therapeutically targeted by novel molecularly targeted agents. Unfortunately, data on replication stress is largely based on experimental models. Further investigation of replication stress in clinical samples is required to optimally implement novel therapeutics. To uncover the relation between oncogene expression, replication stress and clinical features of breast cancer subtypes, we immunohistochemically analyzed the expression of a panel of oncogenes (Cdc25a, Cyclin E and c-Myc) and markers of replication stress (phospho-Ser33-RPA32 and γ-H2AX) in treatment-naive breast tumor tissues (n=384). Triple-negative breast cancers (TNBCs) exhibited the highest levels of phospho-Ser33-RPA32 (*P*<0.001 for all tests) and γ-H2AX (*P*<0.05 for all tests). Moreover, expression levels of Cyclin E (*P*<0.001 for all tests) and c-Myc (*P*<0.001 for all tests) were highest in TNBCs. Expression of Cyclin E positively correlated with phospho-RPA32 (Spearman correlation r=0.37, *P*<0.001) and γ-H2AX (Spearman correlation r=0.63, *P*<0.001). Combined, these data indicate that replication stress is predominantly observed in TNBCs, and is associated with expression levels of Cyclin E. These results indicate that Cyclin E overexpression may be used as a biomarker for patient selection in the clinical evaluation of drugs that target the DNA replication stress response.

## Introduction

Breast cancers are the most frequently diagnosed neoplasms worldwide, with approximately 1.38 million women being diagnosed with breast cancer worldwide every year. One third of these women subsequently die of this disease, accounting for ~14% of all cancer-related deaths in women^1^. Therefore, there is an urgent clinical need for improved breast cancer treatment.

Breast cancers are very heterogeneous, and multiple classification methods have been developed to stratify patient groups. Using gene expression profiling, at least six major breast cancer subtypes have been defined, including ‘normal-like’, ‘luminal A’, ‘luminal B’, ‘HER2-enriched’, ‘claudin-low’ and ‘basal-like’^2^. Furthermore, combining copy number variations with gene expression analysis allowed identification of ten clusters that are associated with differential clinical outcome^3^.

Although multiple classification systems are used in a research context, in standard care breast cancers are subtyped based on the expression of the estrogen and progesterone receptors (ER, PR) and the human epidermal growth factor receptor 2 (HER2), as these receptors are ‘oncogenic drivers’ and relevant drug targets. Breast cancers that do not express the ER, PR or HER2–triple-negative breast cancers (TNBCs) – show a large degree of overlap with the intrinsic ‘basal-like’ and ‘claudin-low’ subtypes, and lack common ‘druggable’ aberrations. TNBC patients do not benefit from anti-hormonal or anti-HER2-targeted treatments, and rely on conventional chemotherapeutic regimens. These tumors display aggressive behavior, and account for ~15-20% of all invasive breast cancers^4^. Although initially high response rate to conventional chemotherapeutics are seen in TNBC, tumors often recur and women have a poor prognosis overall^4^.

Recent genomic analyses of breast cancers have not only underscored the absence of ‘druggable’ oncogenic drivers in TNBC, but also showed that these cancers share a profound genomic instability^5^. This phenomenon is characterized by continuous gains and losses of chromosomal fragments. As a consequence, genomic instability underlies the rapid acquisition of genomic aberrations that drive therapy failure^6^. Finding novel treatment options for genomically instable cancers is not only relevant for TNBCs, but also for other hard-to-treat cancers with extensive genomic instability, including ovarian and pancreatic cancers.

Evidence is increasingly pointing to replication stress as the driver of genomic instability^7,8^. During S-phase of the cell cycle, all DNA must be replicated in a coordinated manner to divide two complete copies of the genome between the two daughter cells. To replicate the entire genome in an orchestrated fashion, replication is initiated at certain genomic loci called ‘replication origins’^8^. Replication origins are not fired simultaneously, but in a temporally controlled way. This sequential firing of the replication origins prevents exhaustion of the nucleotide pool and warrants the availability of essential components of the replication machinery as well as replication checkpoint proteins.

DNA replication can be frustrated in various ways, which is referred to as replication stress. Multiple factors that cause replication stress have been identified, including secondary DNA structures or DNA-RNA hybrids at sites of ongoing transcription^7^. However, the most important source of replication stress appears to be the uncoordinated firing of replication origins due to oncogene activation^7,8,9,10^. As a consequence, oncogene activation depletes the nucleotide pool, leading to slowing or complete stalling of replication forks^11^.

A relevant oncogene related to replication stress is c-Myc, which functions as a transcription factor^12^. c-Myc overexpression promotes cell growth^13^, elevates levels of reactive oxygen species^12^ and causes cellular senescence in experimental models^14^. Another oncogene that has been identified to cause replication stress is Cyclin E. Overexpression of Cyclin E increases the activity of its binding partner Cyclin-dependent Kinase-2 (CDK2) and accelerates S-phase entry. Simultaneously, Cyclin E overexpression results in aberrant activation of origin firing and a consequent nucleotide pool depletion causing replication fork stalling and genomic instability^11,15^. Likewise, the Cdc25a phosphatase activates CDKs, including CDK2, and promotes premature cell cycle progression and checkpoint bypass^16,17,18^.

Cells are equipped with multiple mechanisms to survive replication stress. During replication fork stalling, single-stranded DNA (ssDNA) is exposed and rapidly activates the so-called ‘replication checkpoint’, in which the ATR kinase is the central player^19^. Activation of the replication checkpoint facilitates the rapid coating of ssDNA at replication forks by filaments of replication-Protein A (RPA), which in this process are phosphorylated by ATR^20,21^. RPA filaments subsequently protect and stabilize the otherwise fragile replication forks from nuclease-mediated degradation. When these stalled replication forks are not resolved in time, they can collapse and cause DNA double-strand breaks (DSBs). DSBs rapidly activate the ATM kinase, which leads to activation of multiple downstream targets, including components of the cell cycle machinery and DNA repair pathways^21^. An important ATM substrate at DSBs, including collapsed replication forks, is the histone variant H2AX, which after phosphorylation at serine 139 is referred to as γ-H2AX^22^.

To mitigate the negative impact of replication stress on cellular viability, cancer cells have evolved replication stress-resolving mechanisms. Genomically instable tumors increasingly rely on these mechanisms, including cell cycle checkpoint responses, for their survival^23^. Therefore, components of these replication stress-resolving mechanisms, including the checkpoint kinases Wee1, ATR and Chk1, present promising targets for tumors with high levels of replication stress. Many of the data on replication stress in cancers were derived from experimental model systems. In order to implement novel therapeutic agents that target tumors with high levels of replication stress optimally, it is essential that an inventory is made of tumor subtypes that display replication stress. To this end, and to find potential biomarkers for tumors with high levels of replication stress, we examined replication stress levels in relation to oncogene expression and clinicopathological data in a consecutive well-defined series of breast cancer samples.

## Results

### Patient characteristics

The study population comprised 384 breast cancer patients (Fig. 1a), whose baseline clinical, pathological and treatment characteristics are summarized in Table 1. Breast cancer patients were divided into four subtypes according to their hormone receptor status and HER2 expression. Molecular subtype analysis showed that our cohort consisted of n=161 ER|PR^+^HER2^-^, n=90 ER|PR^+^HER2^+^, n=27 ER|PR^-^ER2^+^ and n=106 ER|PR^-^ER2^-^ (TNBC) patients (Table 1). The age at diagnosis of TNBC patients was lower compared to other patient subgroups (Table 2, *P*<0.05 for all tests), followed by ER|PR^-^ER2^+^ patients^28^. In addition, tumor grade significantly varied across breast cancer subtypes (Table 1, *P*=1.72*10^-13^), and was highest in patients with TNBC (Table 2, *P*<0.05 for all tests). In accordance with treatment guidelines, chemotherapy was most frequently used in TNBC patients (66.0%), but no significant differences between patient subgroups were found (Table 1, *P*=0.08 for all tests). Conversely, radiotherapy and endocrine therapy were predominantly used in breast cancer patients with expression of ER|PR and/or HER2, (Table 2, *P*<0.001 for all tests).

**Figure 1.**
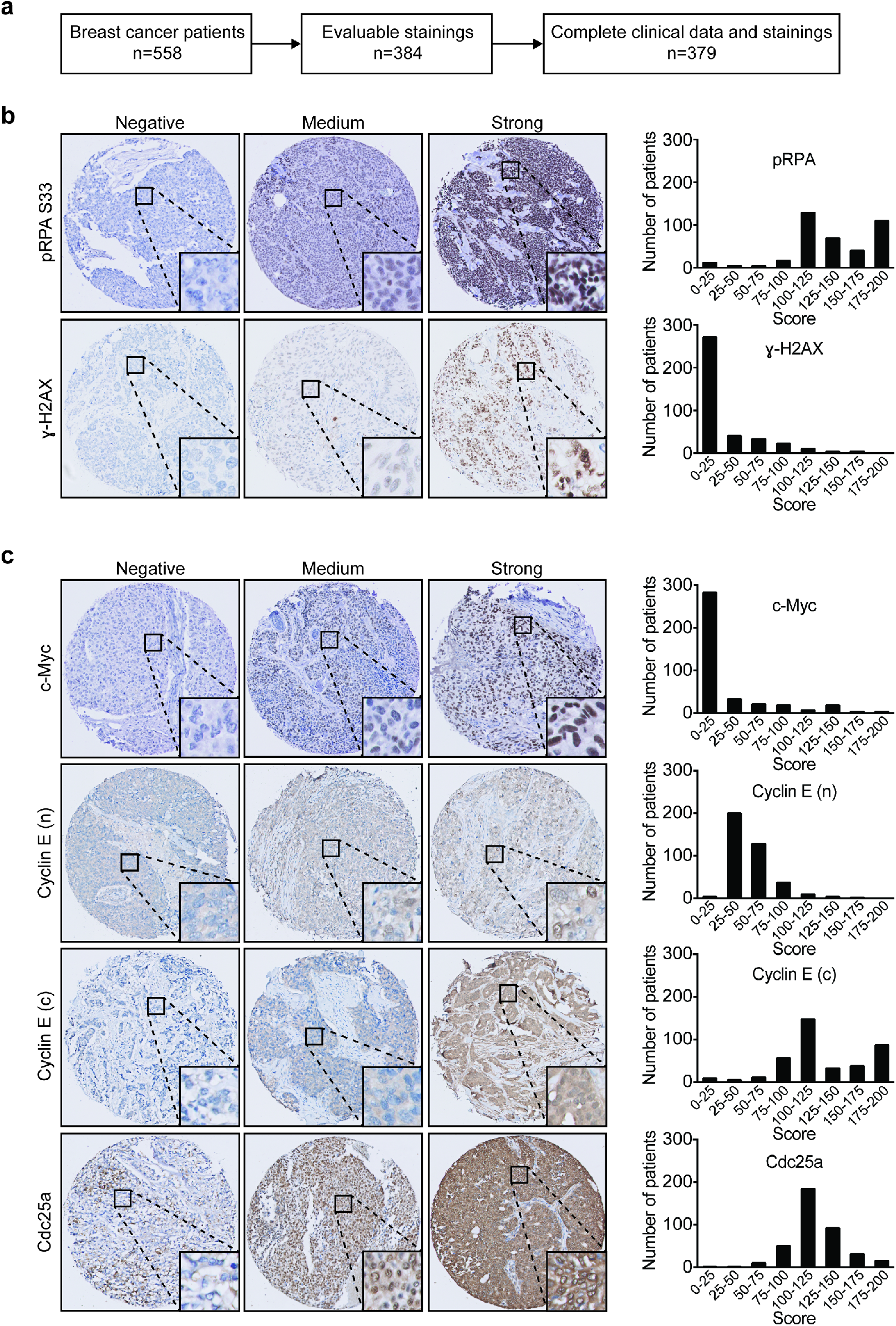
Overview of representative immunohistochemical stainings in breast cancer tissues. **(a)** Flow diagram indicating selection of patients included for immunohistochemical and clinicopathological analyses. **(b)** Analysis of staining intensities of replication stress markers, γ-H2AX and pRPA32 (Ser33) in breast cancer TMAs (n=384). **(c)** Analysis of staining intensities of oncogenes (n=384), including Cdc25a, cytoplasmic (c) and nuclear (n) Cyclin E, and c-Myc.

**Table 1.** Clinicopathological characteristics of the study population. Overview of baseline clinical, pathological and treatment characteristics of patients from the study population cohort (n=384) and breast cancer molecular subtypes ER|PR^+^HER2^-^ (n=161), ER|PR^+^HER2^+^ (n=90), ER|PR^-^ER2^+^ (n=27) and ER|PR^-^ER2’(n=106). The indicated *P*-values for tumor stage, tumor grade, radiation therapy, chemotherapy and endocrine therapy were obtained using Pearson Chi-Square tests, whereas the difference in age was assessed using a Kruskal-Wallis test.

| Variable | No. of patients (%) |  |  |  |  | P-value |
| --- | --- | --- | --- | --- | --- | --- |
|  | Combined cohort<br>n=384 | ER\|PR<sup>+</sup><br>HER2<sup>-</sup><br>n=161 | ER\|PR<sup>+</sup><br>HER2<sup>+</sup><br>n=90 | ER\|PR<sup>-</sup><br>HER2<sup>+</sup><br>n=27 | ER\|PR<sup>-</sup><br>HER2<sup>-</sup><br>n=106 |  |
| Age (years) |  |  |  |  |  | 0.036 |
| Median | 56.1 | 56.7 | 58.1 | 56.1 | 52.7 |  |
| Range | 26.6-90.5 | 27.0-88.6 | 35.9-89.3 | 31.4-74.7 | 26.6-90.5 |  |
| T stage |  |  |  |  |  | 0.040 |
| T1 | 210 (54.7) | 101 (62.7) | 43 (47.8) | 13 (48.1) | 53 (50.0) |  |
| T2 | 147 (38.3) | 51 (31.7) | 41 (45.6) | 13 (48.1) | 42 (39.6) |  |
| T3 | 24 (6.3) | 7 (4.3) | 6 (6.7) | 0 (0.0) | 11 (10.4) |  |
| NA | 3 (0.8) | 2 (1.2) | 0 (0.0) | 1 (3.7) | 0 (0.0) |  |
| Tumor grade |  |  |  |  |  | 1.72*10<sup>-13</sup> |
| I | 70 (18.2) | 44 (27.3) | 17 (18.9) | 4 (14.8) | 5 (4.7) |  |
| II | 143 (37.2) | 77 (47.8) | 38 (42.2) | 10 (37.0) | 18 (17.0) |  |
| III | 169 (44.0) | 40 (24.8) | 35 (38.9) | 13 (48.1) | 81 (76.4) |  |
| Unknown | 2 (0.5) | 0 (0.0) | 0 (0.0) | 0 (0.0) | 2 (1.9) |  |
| Radiation therapy |  |  |  |  |  |  |
| No | 174 (45.3) | 58 (36.0) | 41 (45.6) | 7 (25.9) | 68 (64.2) | 1.60*10<sup>-5</sup> |
| Yes | 210 (54.7) | 103 (64.0) | 49 (54.4) | 20 (74.1) | 38 (35.8) |  |
| Chemotherapy |  |  |  |  |  | 0.079 |
| No | 167 (43.5) | 79 (49.1) | 42 (46.7) | 10 (37.0) | 36 (34.0) |  |
| Yes | 217 (56.5) | 82 (50.9) | 48 (53.3) | 17 (63.0) | 70 (66.0) |  |
| Endocrine therapy |  |  |  |  |  | 3.58*10<sup>-10</sup> |
| No | 268 (69.8) | 93 (57.8) | 55 (61.1) | 19 (70.4) | 101 (95.3) |  |
| Yes | 116 (30.2) | 68 (42.2) | 35 (38.9) | 8 (29.6) | 5 (4.7) |  |

### Levels of replication stress in breast cancer subtypes

To determine levels of replication stress, we immunohistochemically examined the expression of pRPA and γ-H2AX in treatment-naive breast cancer tissues. Representative immunohistochemical pRPA and γ-H2AX stainings are shown in Fig. 1b. Among the breast cancer subtypes, the highest pRPA staining scores were found in TNBC tumors (Table 3a, Fig. 2a). Moreover, TNBC tumors also showed higher pRPA levels than non-TNBC tumors (Fig. 2b, *P*=3.79*10^-9^). In addition, the γ-H2AX scores from TNBC tumors were significantly higher than in ER|PR^+^HER2^-^ and ER|PR^+^HER2^+^ subgroups (Table 3a, *P*<0.05 for all tests), followed by ER|PR^-^ER2^+^ tumors (Fig. 2a). Also, TNBC tumors displayed higher levels of γ-H2AX expression when compared to non-TNBCs (Fig. 2b, *P*=3.42*10^-4^). These data show that the expression of replication stress markers varies among breast cancer subtypes, with TNBCs and ER|PR^-^ER2^+^ tumors exhibiting the highest levels.

**Figure 2.**
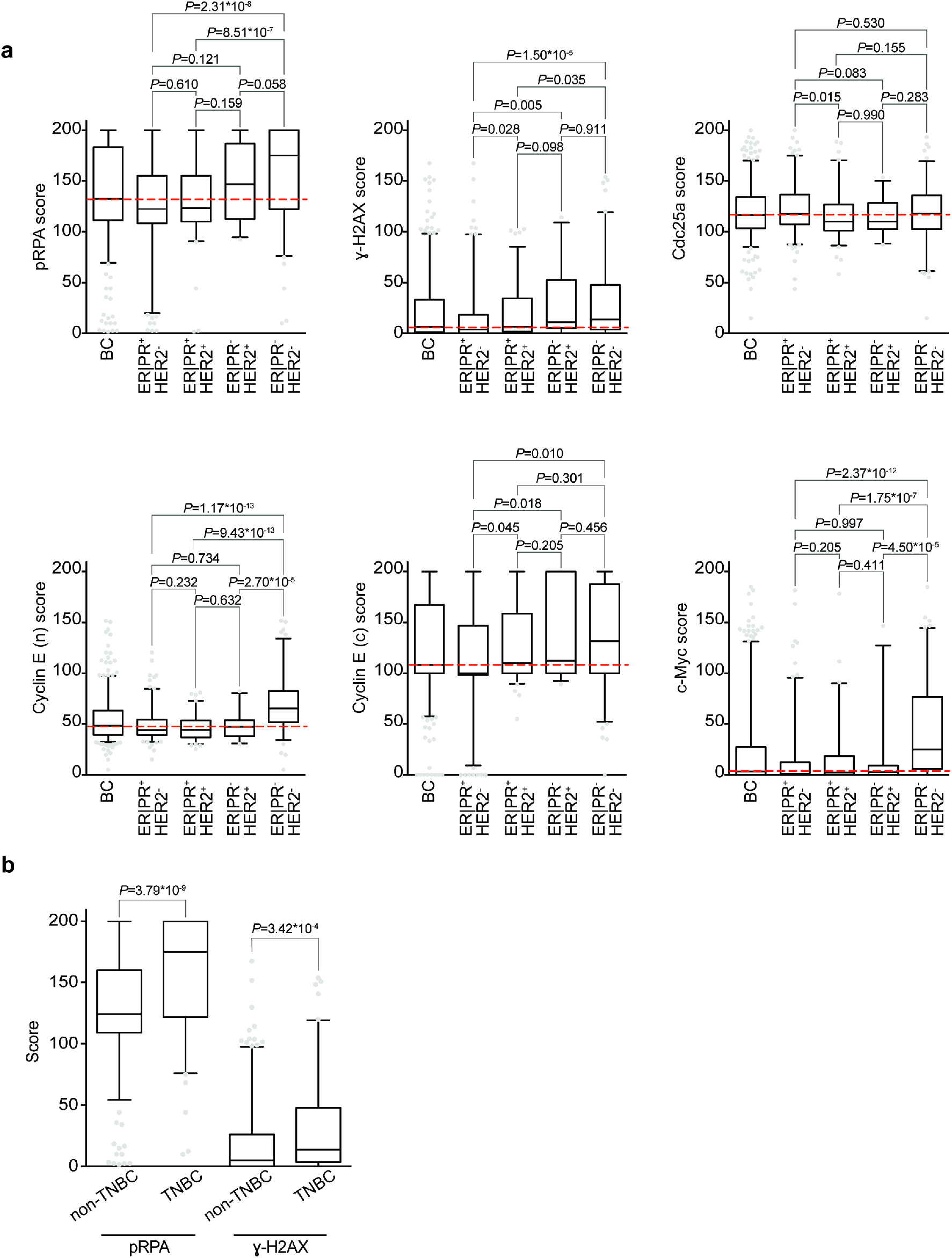
Expression of oncogenes and replication stress markers in breast cancer. **(a)** Patients from the combined cohort (n=384) and breast cancer subtypes ER|PR^+^HER2^-^ (n=161), ER|PR^+^HER2^+^ (n=90), ER|PR^-^ER2^+^ (n=27) and ER|PR^-^ER2^-^(n=106) were analyzed. Tumor tissue was immunohistochemically scored for expression of replication stress markers (pRPA and γ-H2AX) and oncogenes (Cdc25a, Cyclin E (n), Cyclin E (c), c-Myc), indicated *P*-values were calculated using Mann-Whitney U test. **(b)** TNBC patients (ER|PR^-^ER2^-^, n=106) were compared with non-TNBC breast cancers (n=278) for expression of replication stress markers pRPA and γ-H2AX.

### Expression of Cdc25a, Cyclin E and c-Myc in breast cancer subtypes

We next examined the expression levels of Cyclin E (encoded by *CCNE1*) and c-Myc (encoded by *MYC*), two oncogenes that are frequently amplified in TNBC, and have been associated with replication stress in experimental models^29,30,31,32^. For Cyclin E, we separately assessed nuclear and cytoplasmic Cyclin E (Fig. 1c), since cytoplasmic Cyclin E has been related to reduced breast cancer survival^27,33^. We also assessed expression of the Cdc25a phosphatase (Fig. 1c). Although Cdc25a is less frequently overexpressed in breast cancer^16^, Cdc25a overexpression is frequently used to induce replication stress in experimental models^34^, and has been linked to oncogenic activity^35,36^, and for that reason was included in our analysis.

Expression levels of nuclear Cyclin E were significantly higher in TNBC than in the other breast cancer subtypes (Fig. 2a, Table 3b, *P*<0.05 for all tests). In contrast, cytoplasmic Cyclin E levels were high in both TNBC and ER|PR^-^ER2^+^ tumors, compared to ER|PR^+^HER2^-^ tumors (Fig. 2a, Table 3b, *P*<0.05 for all tests). Expression levels of c-Myc were also higher in TNBC (Fig. 2a, Table 3b, *P*<0.001) compared to the other subgroups. Even though TNBC tumors also displayed the highest levels of Cdc25a (Table 3b), these differences were not statistically significant (Fig. 2a and Table 3b). These findings show that TNBCs displayed the highest levels of nuclear Cyclin E and c-Myc. In contrast, Cdc25a expression was similarly distributed among breast cancer subtypes.

Since the highest expression levels of c-Myc and Cyclin E as well as replication stress markers were observed in TNBC, we further analyzed relevant TNBC subgroups. Expression of the androgen receptor (AR) has been described to define a TNBC subgroup with distinct characteristics^37^. In our cohort, n=29 out of 106 TNBC cases (27.4%) expressed the AR (Fig. S1a, Table S2). We next analyzed the expression levels of replication stress markers in TNBC-AR^-^ and TNBC-AR+ subgroups (Fig. S1b). Tumor expression of pRPA (Fig. S1b, Table S2a, *P*=0.686) and γ-H2AX (Fig. S1b, Table S2a, *P*=0.798) were not statistically different between the AR+ versus AR^-^ TNBC cases. Likewise, tumor expression of oncogenes was similarly distributed in TNBC-AR^-^ and TNBC-AR+ subgroups (Fig. S1b, Table S2b, P>0.05 for all tests). These data indicate that in this cohort, expression of replication stress markers and expression of oncogenes are similarly distributed in TNBC-AR^-^ and TNBC-AR+ subgroups.

### Correlations between replication stress markers and Cdc25a, Cyclin E and c-Myc expression

To determine whether expression of Cyclin E, c-Myc or Cdc25a was associated with replication stress in our study population, we examined associations between expression of Cdc25a, Cyclin E and c-Myc with expression of replication stress markers pRPA and γ-H2AX (Table 2). Expression levels of pRPA were positively correlated with those of c-Myc (Table 4a, r=0.26, *P*<0.001), as well as expression levels of nuclear Cyclin E (Table 4a, r=0.37, *P*<0.001) and cytoplasmic Cyclin E (Table 4a, r=0.28, *P*<0.001) in the entire cohort. Among breast cancer subtypes, the strongest correlations were found in TNBC between Cyclin E and pRPA expression (Table 4a, r=0.43, *P*<0.001), and between c-Myc and pRPA (Table 4a, r=0.36, *P*<0.001). Furthermore, Cyclin E expression was strongly correlated with levels of γ-H2AX staining (Table 4b, r=0.63, *P*<0.001). Spearman correlation analysis of breast cancer subtypes revealed that the association between Cyclin E and γ-H2AX expression was strongest in ER|PR^-^ER2^+^ (Table 4b, r=0.86, *P*<0.001) and TNBC (Table 4b, r=0.71, *P*<0.001). Combined, these data indicate that expression of Cyclin E is associated with expression of replication stress markers in our study population, especially in the TNBC and ER|PR^-^ER2^+^ subgroups.

**Table 2.** Relation between breast cancer subtypes and clinicopathological characteristics in the study population. The indicated *P*-values were obtained using a Pearson Chi-Square test. The *P*-values for age were assessed using a Mann-Whitney U test.

| BC subtype | Patient characteristics: *P*-value |  |  |  |  |  |  |  |  |
| --- | --- | --- | --- | --- | --- | --- | --- | --- | --- |
|  | ER\|PR<sup>+</sup>HER2<sup>+</sup> |  |  | ER\|PR<sup>-</sup>HER2<sup>-</sup> |  |  | ER\|PR<sup>-</sup>HER2<sup>+</sup> |  |  |
|  | Age | T-Stage | T-Grade | Age | T-Stage | T-Grade | Age | T-Stage | T-Grade |
| ER\|PR<sup>+</sup>HER2<sup>-</sup> | 0.358 | 0.072 | 0.052 | 0.029 | 0.052 | 2.91*10<sup>-16</sup> | 0.434 | 0.195 | 0.040 |
| ER\|PR<sup>+</sup>HER2<sup>+</sup> |  |  |  | 0.007 | 0.542 | 3.48*10<sup>-7</sup> | 0.244 | 0.160 | 0.683 |
| ER\|PR<sup>-</sup>HER2<sup>-</sup> |  |  |  |  |  |  | 0.411 | 0.068 | 0.016 |
| BC subtype | Treatment: *P*-value |  |  |  |  |  |  |  |  |
|  | ER\|PR<sup>+</sup>HER2<sup>+</sup> |  |  | ER\|PR<sup>-</sup>HER2<sup>-</sup> |  |  | ER\|PR<sup>-</sup>HER2<sup>+</sup> |  |  |
|  | Chemo | Radio | Endo | Chemo | Radio | Endo | Chemo | Radio | Endo |
| ER\|PR<sup>+</sup>HER2<sup>-</sup> | 0.715 | 0.138 | 0.605 | 0.015 | 7.00*10<sup>-6</sup> | 1.70*10<sup>-11</sup> | 0.247 | 0.307 | 0.217 |
| ER\|PR<sup>+</sup>HER2<sup>+</sup> |  |  |  | 0.070 | 0.009 | 3.31*10<sup>-9</sup> | 0.377 | 0.069 | 0.381 |
| ER\|PR<sup>-</sup>HER2<sup>-</sup> |  |  |  |  |  |  | 0.764 | 3.49*10<sup>-4</sup> | 1.00*10<sup>-4</sup> |

**Table 3a.** Tumor expression of replication stress markers and oncogenes in the study population. **(a)** Tumor expression of pRPA and γ-H2AX in the combined cohort (n=384) and breast cancer subtypes ER|PR^+^HER2^-^ (n=161), ER|PR^+^HER2^+^ (n=90), ER|PR^-^ER2^+^ (n=27) and ER|PR^-^ER2’(n=106).

| BC subtype | TMA score |  |  |  |  |
| --- | --- | --- | --- | --- | --- |
|  | Mean | SD | P5 | P50 | P95 |
| pRPA |  |  |  |  |  |
| Combined cohort (n=384) | 139.1 | 44.7 | 69.4 | 132.5 | 200.0 |
| ER\|PR<sup>+</sup>HER2<sup>-</sup> (n=161) | 128.2 | 45.0 | 19.9 | 122.5 | 200.0 |
| ER\|PR<sup>+</sup>HER2<sup>+</sup> (n=90) | 132.0 | 38.0 | 90.7 | 123.3 | 200.0 |
| ER\|PR<sup>-</sup>HER2<sup>+</sup> (n=27) | 147.8 | 38.5 | 94.5 | 146.7 | 200.0 |
| ER\|PR<sup>-</sup>HER2<sup>-</sup> (n=106) | 159.4 | 44.3 | 76.2 | 175.0 | 200.0 |
| γ-H2AX | | | | | |
| Combined cohort (n=384) | 23.1 | 33.7 | 0.0 | 6.3 | 98.3 |
| ER\|PR<sup>+</sup>HER2<sup>-</sup> (n=161) | 18.5 | 32.7 | 0.0 | 3.8 | 97.4 |
| ER\|PR<sup>+</sup>HER2<sup>+</sup> (n=90) | 20.6 | 27.4 | 0.0 | 6.5 | 85.5 |
| ER\|PR<sup>-</sup>HER2<sup>+</sup> (n=27) | 30.4 | 35.6 | 0.0 | 10.8 | 109.2 |
| ER\|PR<sup>-</sup>HER2<sup>-</sup> (n=106) | 30.6 | 38.3 | 0.0 | 13.6 | 119.1 |

**Table 3b.** **(b)** Tumor expression of Cdc25a, Cyclin E (n), Cyclin E (c) and c-Myc in the combined cohort (n=384) and breast cancer subtypes ER|PR^+^HER2^-^ (n=161), ER|PR^+^HER2^+^ (n=90), ER|PR^-^ER2^+^ (n=27) and ER|PR^-^ER2’(n=106).

| BC subtype | TMA score |  |  |  |  |
| --- | --- | --- | --- | --- | --- |
|  | Mean | SD | P5 | P50 | P95 |
| Cdc25a |  |  |  |  |  |
| Combined cohort (n=384) | 119.5 | 25.7 | 85.0 | 116.7 | 170.0 |
| ER\|PR<sup>+</sup>HER2<sup>-</sup> (n=161) | 122.4 | 25.1 | 87.6 | 117.5 | 174.9 |
| ER\|PR<sup>+</sup>HER2<sup>+</sup> (n=90) | 116.5 | 24.5 | 86.4 | 110.0 | 170.4 |
| ER\|PR<sup>-</sup>HER2<sup>+</sup> (n=27) | 115.0 | 18.0 | 88.5 | 110.0 | 150.0 |
| ER\|PR<sup>-</sup>HER2<sup>-</sup> (n=106) | 118.8 | 28.9 | 61.4 | 117.9 | 169.6 |
| Cyclin E (n) |  |  |  |  |  |
| Combined cohort (n=384) | 54.1 | 21.4 | 32.3 | 48.3 | 97.5 |
| ER\|PR<sup>+</sup>HER2<sup>-</sup> (n=161) | 49.1 | 16.1 | 32.6 | 43.9 | 84.8 |
| ER\|PR<sup>+</sup>HER2<sup>+</sup> (n=90) | 46.5 | 12.7 | 30.4 | 44.1 | 72.9 |
| ER\|PR<sup>-</sup>HER2<sup>+</sup> (n=27) | 48.5 | 14.8 | 30.9 | 47.2 | 80.5 |
| ER\|PR<sup>-</sup>HER2<sup>-</sup> (n=106) | 69.4 | 27.5 | 34.2 | 65.3 | 134.3 |
| Cyclin E (c) |  |  |  |  |  |
| Combined cohort (n=384) | 126.4 | 46.7 | 57.5 | 108.3 | 200.0 |
| ER\|PR<sup>+</sup>HER2<sup>-</sup> (n=161) | 117.3 | 48.9 | 9.3 | 100.0 | 200.0 |
| ER\|PR<sup>+</sup>HER2<sup>+</sup> (n=90) | 128.0 | 37.7 | 89.6 | 110.0 | 200.0 |
| ER\|PR<sup>-</sup>HER2<sup>+</sup> (n=27) | 140.5 | 44.1 | 92.5 | 112.5 | 200.0 |
| ER\|PR<sup>-</sup>HER2<sup>-</sup> (n=106) | 135.3 | 48.6 | 52.3 | 131.7 | 200.0 |
| c-Myc |  |  |  |  |  |
| Combined cohort (n=384) | 24.1 | 40.4 | 0.0 | 3.3 | 131.3 |
| ER\|PR<sup>+</sup>HER2<sup>-</sup> (n=161) | 15.1 | 32.6 | 0.0 | 0.8 | 95.7 |
| ER\|PR<sup>+</sup>HER2<sup>+</sup> (n=90) | 16.6 | 30.2 | 0.0 | 2.1 | 90.0 |
| ER\|PR<sup>-</sup>HER2<sup>+</sup> (n=27) | 14.7 | 33.9 | 0.0 | 2.5 | 127.3 |
| ER\|PR<sup>-</sup>HER2<sup>-</sup> (n=106) | 46.5 | 50.8 | 0.0 | 25.0 | 144.4 |

**Table 4a.** Spearman rank correlation analysis of replication stress marker expression versus oncogene expression in the study population. **(a)** Association analysis between oncogene expression and pRPA expression in the combined cohort (n=384) and in breast cancer subtypes.

| Variable | BC Subtype |  |  |  |  |
| --- | --- | --- | --- | --- | --- |
|  | Combined cohort | ER\|PR<sup>+</sup><br>HER2<sup>-</sup> | ER\|PR<sup>+</sup><br>HER2<sup>+</sup> | ER\|PR<sup>-</sup><br>HER2<sup>+</sup> | ER\|PR<sup>-</sup><br>HER2<sup>-</sup> |
|  | n=384 | n=161 | n=90 | n=27 | n=106 |
| pRPA |  |  |  |  |  |
| Cdc25a |  |  |  |  |  |
| Correlation | -0.135 | -0.269 | -0.152 | 0.076 | 0.035 |
| P-value | 0.008 | 0.001 | 0.153 | 0.707 | 0.719 |
| Cyclin E (n) |  |  |  |  |  |
| Correlation | 0.371 | 0.090 | 0.289 | 0.232 | 0.432 |
| P-value | 5.61*10<sup>-14</sup> | 0.258 | 0.006 | 0.245 | 4.00*10<sup>-6</sup> |
| CyclinE (c) |  |  |  |  |  |
| Correlation | 0.275 | 0.309 | 0.143 | 0.218 | 0.262 |
| P-value | 4.25*10<sup>-8</sup> | 6.50*10<sup>-5</sup> | 0.180 | 0.274 | 0.007 |
| c-Myc |  |  |  |  |  |
| Correlation | 0.256 | 0.214 | 0.180 | 0.133 | 0.359 |
| P-value | 3.59*10<sup>-7</sup> | 0.006 | 0.090 | 0.509 | 1.58*10<sup>-4</sup> |

**Table 4b.** **(b)** Association analysis between oncogene expression and γ-H2AX expression in the combined cohort (n=384) and in breast cancer subtypes.

| Variable | BC Subtype |  |  |  |  |
| --- | --- | --- | --- | --- | --- |
|  | Combined cohort | ER\|PR<sup>+</sup><br>HER2<sup>-</sup> | ER\|PR<sup>+</sup><br>HER2<sup>+</sup> | ER\|PR<sup>-</sup><br>HER2<sup>+</sup> | ER\|PR<sup>-</sup><br>HER2<sup>-</sup> |
|  | n=384 | n=161 | n=90 | n=27 | n=106 |
| γ-H2AX |  |  |  |  |  |
| Cdc25a |  |  |  |  |  |
| Correlation | -0.030 | -0.140 | -0.037 | 0.368 | 0.170 |
| P-value | 0.561 | 0.076 | 0.732 | 0.059 | 0.081 |
| Cyclin E (n) |  |  |  |  |  |
| Correlation | 0.632 | 0.538 | 0.613 | 0.855 | 0.705 |
| P-value | 3.07*10<sup>-44</sup> | 1.80*10<sup>-13</sup> | 1.34*10<sup>-10</sup> | 1.40*10<sup>-8</sup> | 3.53*10<sup>-17</sup> |
| CyclinE (c) |  |  |  |  |  |
| Correlation | 0.192 | 0.209 | 0.258 | -0.260 | 0.078 |
| P-value | 1.55*10<sup>-4</sup> | 0.008 | 0.014 | 0.190 | 0.424 |
| c-Myc |  |  |  |  |  |
| Correlation | 0.129 | 0.113 | 0.032 | 0.214 | -0.032 |
| P-value | 0.011 | 0.153 | 0.763 | 0.283 | 0.743 |

To test whether the association between oncogene expression and expression of replication stress markers was affected by AR expression in TNBCs, Spearman correlation analyses were performed. A stronger association between pRPA and Cyclin E was observed in AR^-^ negative TNBCs (Table S3a, r=0.463, *P*<0.001), when compared to AR-positive TNBCs (Table S3a, r=0.327, *P*=0.083). Similarly, Spearman correlation analysis underscored that the association between Cyclin E and γ-H2AX was stronger in AR-negative tumors (Table S3b, r=0.755, *P*<0.05) than in AR-positive tumors (Table S3b, r=0.574, *P*=0.001). In line with this observation, Cdc25a and Cyclin E showed weaker associations in AR-positive tumors (Table S3a, Cdc25a: r=-0.214, *P*=0.265; Cyclin E: r=0.092, *P*=0.637), than in AR-negative tumors (Table S3a, Cdc25a: r=0.131, *P*=0.256; Cyclin E: r=0.320, *P*=0.005). In addition, AR-negative tumors showed stronger associations between γ-H2AX and Cdc25a (Table S3b, r=0.275, *P*=0.016) and between γ-H2AX and Cyclin E (Table S3b, r=0.118, *P*=0.306) than AR-positive tumors (Table S3b, Cdc25a: r=-0.100, *P*=0.607 and (Cyclin E: r=0.048, *P*=0.806). In conclusion, markers of replication stress appear equally expressed in AR-negative and AR-positive TNBCs, although the associations between replication stress (pRPA, γ-H2AX) and oncogene expression (Cdc25a, Cyclin E) are strongest in AR-negative TNBCs within our cohort.

### Associations of expression of replication stress markers with clinicopathological characteristics and tumor expression of Cdc25a, Cyclin E and c-Myc

Linear regression analyses were performed to evaluate the relation between expression of replication stress markers versus clinicopathological characteristics and tumor expression of Cdc25a, Cyclin E and c-Myc. Univariate regression analysis showed that pRPA was associated with γ-H2AX (Table 5, β=0.409, *P*<0.001). In contrast, oncogenes presented weak associations with pRPA (Table 5). The covariates from univariate regression analyses that displayed *P*<0.05 were included for multivariate analyses, which showed that pRPA was weakly related to γ-H2AX (Table 5, β=0.351, *P*<0.001). Conversely, tumor expression of γ-H2AX was associated with nuclear Cyclin E (Table 6, β=0.765, *P*<0.001). No significant associations were found between clinicopathological parameters and pRPA or γ-H2AX expression.

**Table 5.** Relation between pRPA expression versus oncogene expression and clinicopathological characteristics of the study population. Univariate linear regression analyses were performed to analyze the relation between expression of the replication stress marker pRPA versus clinicopathological characteristics and tumor expression of Cdc25a, Cyclin E or c-Myc (n=379). Comparisons with a P value <0.05 in univariate linear regression analyses were selected for multivariate linear regression analyses.

|  | pRPA |  |  |  |  |  |
| --- | --- | --- | --- | --- | --- | --- |
|  | Univariate |  |  | Multivariate |  |  |
|  | Beta | 95% CI | P-value | Beta | 95% CI | P-value |
| Age (continuous) | -0.125 | -0.684-[-0.073] | 0.015 | -0.064 | -0.476-0.089 | 0.178 |
| Tumor subtype |  |  |  |  |  |  |
| ER\|PR<sup>+</sup>HER2<sup>-</sup> | *Ref.* |  |  | *Ref.* |  |  |
| ER\|PR<sup>+</sup>HER2<sup>+</sup> | 0.037 | -7.293-14.989 | 0.497 | 0.024 | -7.663-12.610 | 0.632 |
| ER\|PR<sup>-</sup>HER2<sup>+</sup> | 0.105 | 0.755-36.488 | 0.041 | 0.067 | -4.499-28.145 | 0.155 |
| ER\|PR<sup>-</sup>HER2<sup>-</sup> | 0.304 | 19.817-41.120 | 3.64*10<sup>-8</sup> | 0.188 | 7.038-30.569 | 0.002 |
| Tumor stage |  |  |  |  |  |  |
| I | *Ref.* |  |  |  |  |  |
| II | 0.042 | -5.638-13.349 | 0.425 |  |  |  |
| III | -0.059 | -29.687-8.204 | 0.266 |  |  |  |
| Tumor grade |  |  |  |  |  |  |
| I | *Ref.* |  |  | *Ref.* |  |  |
| II | 0.001 | -12.632-12.902 | 0.983 | -0.019 | -12.912-9.386 | 0.756 |
| III | 0.165 | 2.439-27.318 | 0.019 | -0.098 | -21.113-3.502 | 0.160 |
| Cdc25a | -0.055 | -0.273-0.080 | 0.284 |  |  |  |
| CyclinE (n) | 0.377 | 0.589-0.979 | 3.13*10<sup>-14</sup> | 0.028 | -0.246-0.364 | 0.703 |
| CyclinE (c) | 0.272 | 0.167-0.353 | 7.29*10<sup>-8</sup> | 0.188 | 0.092-0.267 | 6.90*10<sup>-5</sup> |
| c-Myc | 0.248 | 0.167-0.386 | 1.00*10<sup>-6</sup> | 0.150 | 0.055-0.280 | 0.004 |
| γ-H2AX | 0.409 | 0.421-0.667 | 1.03*10<sup>-16</sup> | 0.351 | 0.292-0.643 | 2.70*10<sup>-7</sup> |

**Table 6.**
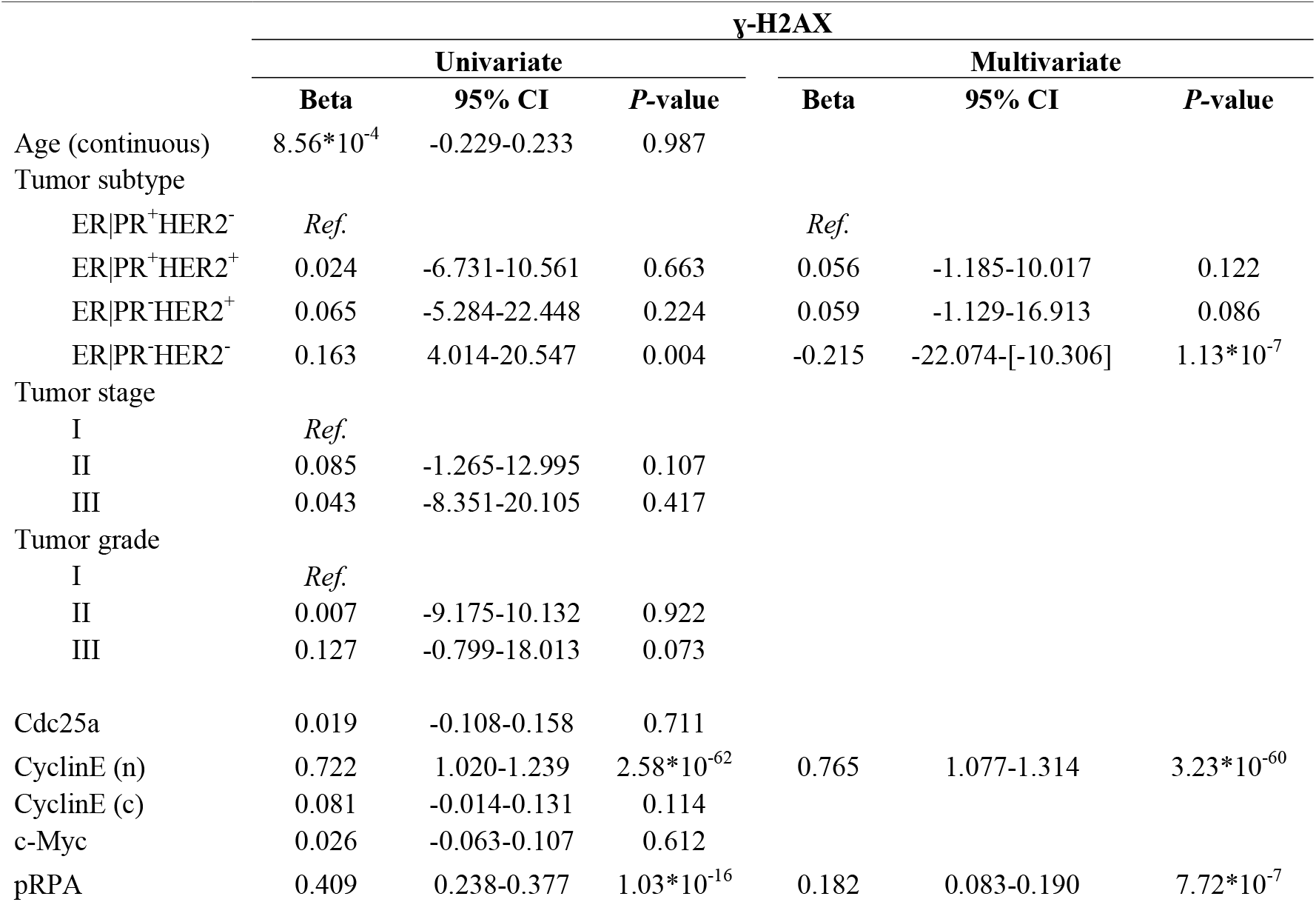
Relation between γ-H2AX expression versus oncogene expression and clinicopathological characteristics of the study population. Univariate linear regression analyses were performed to analyze the relation between expression of the replication stress marker γ-H2AX versus clinicopathological characteristics and tumor expression of Cdc25a, Cyclin E or c-Myc (n=379). Comparisons with a P value <0.05 in univariate linear regression analyses were selected for multivariate linear regression analyses.

| | γ-H2AX | | | | | |
| --- | --- | --- | --- | --- | --- | --- |
|  | Univariate |  |  | Multivariate |  |  |
|  | Beta | 95% CI | P-value | Beta | 95% CI | P-value |
| Age (continuous) | $8.56 \times 10^{-4}$ | -0.229-0.233 | 0.987 | | | |
| Tumor subtype |  |  |  |  |  |  |
| ER\|PR<sup>+</sup>HER2<sup>-</sup> | *Ref.* |  |  | *Ref.* |  |  |
| ER\|PR<sup>+</sup>HER2<sup>+</sup> | 0.024 | -6.731-10.561 | 0.663 | 0.056 | -1.185-10.017 | 0.122 |
| ER\|PR<sup>-</sup>HER2<sup>+</sup> | 0.065 | -5.284-22.448 | 0.224 | 0.059 | -1.129-16.913 | 0.086 |
| ER\|PR<sup>-</sup>HER2<sup>-</sup> | 0.163 | 4.014-20.547 | 0.004 | -0.215 | -22.074-[-10.306] | $1.13 \times 10^{-7}$ |
| Tumor stage |  |  |  |  |  |  |
| I | *Ref.* |  |  |  |  |  |
| II | 0.085 | -1.265-12.995 | 0.107 |  |  |  |
| III | 0.043 | -8.351-20.105 | 0.417 |  |  |  |
| Tumor grade |  |  |  |  |  |  |
| I | *Ref.* |  |  |  |  |  |
| II | 0.007 | -9.175-10.132 | 0.922 |  |  |  |
| III | 0.127 | -0.799-18.013 | 0.073 |  |  |  |
| Cdc25a | 0.019 | -0.108-0.158 | 0.711 |  |  |  |
| CyclinE (n) | 0.722 | 1.020-1.239 | $2.58 \times 10^{-62}$ | 0.765 | 1.077-1.314 | $3.23 \times 10^{-60}$ |
| CyclinE (c) | 0.081 | -0.014-0.131 | 0.114 |  |  |  |
| c-Myc | 0.026 | -0.063-0.107 | 0.612 |  |  |  |
| pRPA | 0.409 | 0.238-0.377 | $1.03 \times 10^{-16}$ | 0.182 | 0.083-0.190 | $7.72 \times 10^{-7}$ |

In summary, these findings indicate that replication stress measured with pRPA is related to γ-H2AX, whereas replication stress quantified with γ-H2AX is positively associated with tumor expression of nuclear Cyclin E.

## Discussion

In the present study, we examined the relation between oncogene expression and replication stress marker expression in breast cancer subtypes. Our immunohistochemical analyses show that levels of replication stress and oncogene expression vary among breast cancer subtypes. Notably, the highest expression levels of replication stress markers and oncogenes were found in the TNBC and ER|PR^-^ER2^+^ subgroups. Furthermore, nuclear Cyclin E expression, and to a lesser extend c-Myc expression, were strongly associated with the levels of replication stress.

Previous immunohistochemical studies showed that Cdc25a was highly expressed in 69.6% of human breast carcinomas analyzed (n=46)^38^. A second study revealed that Cdc25a was overexpressed in 47% of breast cancer cases, in a cohort of 144 patients^16^. In comparison, in the present study 29.5% of tumors displayed strong Cdc25a staining intensity. Cdc25a has also been shown to be overexpressed in other cancer types, including high-grade serous ovarian cancer and head-and-neck squamous cell carcinoma^35,39^, which are characterized by genomic instability^40,41^.

Concerning c-Myc expression, our results indicated that high c-Myc expression was predominantly observed in TNBCs, and that c-Myc expression was associated with expression of the replication stress marker pRPA. These findings are in line with previous reports, showing frequent *MYC* amplification in TNBC^29^. Importantly, our results provide confirmation that the link between c-Myc-overexpression and induction of replication stress is also observed in patient samples ^12,31,42^.

*CCNE1* is frequently amplified in TNBC, in line with our finding that high Cyclin E expression is most prominent in TNBC cases^29^. Importantly, our observation that Cyclin E expression is associated with expression of replication stress markers is in line with experimental models, in which Cyclin E overexpression has been shown to trigger a DNA damage response^11,15,43^. Specific isoforms of Cyclin E, so-called low molecular-weight Cyclin E isoforms (LMW-E) are suggested to accumulate in the cytoplasm because they lack the NH_2_-terminal nuclear localization signal^44^. Although expression of cytoplasmic Cyclin E has previously been shown to induce various features that relate to replication stress, including chromosome missegregation^45^, our data show that cytoplasmic Cyclin E expression is similarly distributed among breast cancer subtypes, and is not associated with expression of replication stress markers pRPA and γ-H2AX.

Taken together, our findings indicate that among breast cancer subtypes, TNBCs and ER|PR^-^HER2^+^ tumors are characterized by overexpression of the c-Myc and Cyclin E oncogenes, and by higher expression levels of replication stress markers. These findings are relevant, as increasing numbers of drugs are being developed that target cancer cells with high levels of replication stress. Specifically, inhibitors of the cell cycle checkpoint kinases Chk1 and ATR are currently being tested in combination with genotoxic agents that interfere with DNA replication^46,47^. In parallel, inhibitors of the Wee1 kinase have been developed. The potential of Wee1 inhibition was early on attributed to high levels of replication stress^48^ and preclinical data indicated that Wee1 inhibition would be preferentially effective in Cyclin E-overexpressing cancer cells^49^. In line with these data, ovarian cancer patients that responded favorably to Wee1 inhibitor treatment more frequently showed tumor overexpression of Cyclin E^50^. Based on these observations, a clinical trial testing Wee1 inhibitor treatment in patients selected on *CCNE1* amplification is currently ongoing (clinicaltrials.gov identifier: NCT03253679).

Although different cell cycle checkpoint inhibitors are already in clinical development, an effective patient selection strategy is required to identify those patients who might benefit from these drugs. For breast cancer patients, our data underscore that overexpression of nuclear Cyclin E could be used as a selection criterion for treatment with drugs that target replication stress, including inhibitors of Wee1, Chk1 and ATR.

## Materials and methods

### Breast cancer tissue

Immunohistochemical analysis was performed on treatment-naive tissue specimens of 558 patients with breast cancer who underwent surgery at the University Medical Center Groningen (UMCG). A first cohort (cohort A) consisted of 450 consecutive patients, of whom tumor specimens were collected between 1996 and 2005, considering the availability of sufficient paraffin-embedded tissue. A second cohort (cohort B) consisted of 108 consecutive patients with TNBC, of whom tumor specimens were collected from 2005 to 2010. Clinicopathological data gathered in this study was anonymized, and used in accordance with the Declaration of Helsinki, and in line with the regulations posed by the Institutional Review Board (IRB) of the UMCG. Tumor specimens were processed to generate nine tissue microarrays (TMAs). The first seven TMAs contained samples from cohort A, as described previously^24,25^. The final two TMAs contained TNBC cases from cohort B. For each tumor, three different cores were included in TMAs.

### Immunohistochemistry

TMA slices were deparaffinized using xylene. Slides were incubated for 30 minutes in 0.3% hydrogen peroxidase (H_2_O_2_) to suppress endogenous peroxidase activity. Immunohistochemistry was performed with primary antibodies against Cdc25a (1:400; rabbit, #sc-97, clone 144; Santa Cruz Biotechnology, CA, USA), Cyclin E (1:1000; rabbit, #sc-198, clone C19; Santa Cruz Biotechnology, CA, USA), c-Myc (RTU; rabbit, #790-4628, clone Y69; Roche, Basel, Switzerland), phospho-RPA32 (Ser33) (1:6400; rabbit, #A300-246A, clone S33; Bethyl, Texas, USA), γ-H2AX (1:300; mouse, #05-636, clone JBW301; Millipore, Amsterdam, The Netherlands) and the androgen receptor (AR) (RTU; rabbit, #760-4605, clone SP107; Roche, Basel, Switzerland). Staining was detected by the application of 3,3-diaminobenzidine (DAB), and hematoxylin as a counterstaining. For c-Myc and pRPA, the complete staining procedure was performed on an autostainer (BenchMark Ultra IHC/ISH, Roche, Basel, Switzerland). Additional information about antibodies and staining protocols is provided in Supplemental Table 1.

Scoring was performed semi-quantitatively by two independent researchers, without knowledge of clinical data, and was supervised by a breast cancer pathologist. Stainings were categorized according to percentages of cells that showed staining and on intensity of staining. Staining intensity was scored in three categories: 0 (negative), 1 (medium), and 2 (high). In order to calculate the score for each core, the percentage of cells in each group was multiplied by their intensity score, resulting in a range from 0 to 200 points. Next, the scores from each case and staining were averaged and considered for analysis.

For Cdc25a, only nuclear staining was considered, in line with a previous study^26^. For Cyclin E, nuclear and cytoplasmic staining were scored individually^27^. In addition, nuclear c-Myc, pRPA and γ-H2AX stainings were evaluated. A concordance of more than 90% was found between observers. Discordant scores were reviewed and adjusted to consensus. The status of ER, PR, HER2 and AR was determined according to the guidelines of the American Society of Clinical Oncology/College of American Pathologists by counting at least 100 cells.

Immunohistochemical stainings were considered evaluable when a tumor core contained at least 10% tumor cells. In addition, tumor stainings were included for analysis when at least 2 out of 3 cores were evaluable. Core loss over 558 cases was on average 15.1% (Cdc25a, n=106 [18.9%]; nuclear Cyclin E, n=74 [13.2%]; cytoplasmic Cyclin E, n=82 [14.6%]; c-Myc, n=68 [12.1%]; pRPA, n=77 [13.8%] and γ-H2AX, n=99 [17.7%]). For 384 out of 558 cases, all five stainings (pRPA, γ-H2AX, Cyclin E, c-Myc and Cdc25a) were evaluable. Of 384 cases with complete evaluable immunohistochemical stainings, 379 had complete clinicopathological data available, and were included for statistical analysis.

### Statistical analyses

Analyses were performed on the total study population as well as on four patient subgroups based on hormone receptor status and HER2 expression. Differences regarding clinicopathological features, treatment, and immunohistochemical expression levels between the four groups were analyzed using Pearson Chi-Square tests in case of categorical variables, while Kruskal-Wallis tests and Mann-Whitney U tests were used in case of continuous variables.

Univariate linear regression analyses were performed to study the relation between expression of replication stress markers versus clinicopathological characteristics and tumor expression of Cdc25a, Cyclin E or c-Myc. Comparisons that reached *P*< 0.05 in univariate linear regression analyses were selected for multivariate linear regression analyses. All statistical analyses in this study were performed using SPSS Statistics 23.0 (IBM).

## Acknowledgements

This work was financially supported by grants from the Netherlands Organization for Scientific research (NWO-VIDI #91713334 to M.A.T.M.v.V.) and the European Research Council (ERC CoS Grant 682421 to M.A.T.M.v.V.).

## Author contributions

S.G.LL., T.d.B., B.v.d.V., and M.A.T.M.v.V. designed the experiments. S.G.LL., E.J. and M.E. performed IHC stainings and scoring. S.G.LL. performed the statistical analyses. M.A.T.M.v.V, B.v.d.V. and T.d.B. supervised the statistical analyses. M.A.T.M.v.V. and S.G.LL. wrote the manuscript with input from all authors.

## Data availability statement

The data that support the findings of this study are available from the corresponding author upon reasonable request.

---

**Supplemental Fig. 1: Androgen receptor expression in triple-negative breast cancers. (a)** Representative immunohistochemical stainings of the androgen receptor in TNBC patients (ER|PR^-^ER2^-^, n=106). **(b)** TNBC patients were subclassified based on the presence or absence of AR staining. TNBC-AR^-^ (n=77) and TNBC-AR+ (n=29) were compared based on their scores on replication stress markers and oncogenes.

**Supplemental Table 1: List of antibodies and protocols for immunohistochemical analysis**. Information regarding antibodies, antigen retrieval methods and detection kits is presented.

**Supplemental Table 2: Tumor expression of replication stress markers and oncogenes in the TNBC-AR subgroups. (a)** Tumor expression of replication stress markers pRPA and γ-H2AX in the combined TNBC cohort (n=106), and the TNBC subtypes ER|PR^-^ HER2^-^AR^-^ (n=77) and ER|PR^-^ER2^+^AR+ (n=29). **(b)** Tumor expression of oncogenes Cdc25a, Cyclin E (n), Cyclin E (c) and c-Myc in the combined TNBC cohort (n=106), and the TNBC subtypes ER|PR^-^ER2^-^AR^-^ (n=77) and ER|PR^-^HER2^+^AR+ (n=29).

**Supplemental Table 3: Spearman rank correlation of replication stress marker expression versus oncogene expression in the TNBC-AR subgroups. (a)** Association analysis between oncogene expression and pRPA expression in the combined TNBC cohort (n=106) and the indicated TNBC subtypes. **(b)** Association analysis between oncogene expression and γ-H2AX expression in the combined TNBC cohort (n=106) and the indicated TNBC subtypes.

## Abbreviations

AR: androgen receptor
CDK2: cyclin-dependent kinase-2
DAB: 3,3-diaminobenzidine
DSBs: double-strand breaks
ER: estrogen receptor
HER2: human epidermal growth factor receptor-2
H_2_O_2_: hydrogen peroxidase
IRB: Institutional Review Board
PR: progesterone receptor
RPA: replication protein-A
ssDNA: single-stranded DNA
TMAs: tissue microarrays
TNBCs: triple-negative breast cancers

